# Engineering Nanobody Conformational Switches

**DOI:** 10.64898/2026.11.06.764871

**Authors:** Alba Pejenaute, Laura Gálvez-Larrosa, Pablo Herrero-Alfonso, Elena Gómez-Rubio, Santos Merino, Óscar Millet, Francesca Peccati, Gonzalo Jiménez-Osés, Gabriel Ortega-Quintanilla

## Abstract

Conformational signaling is widely used in biology, but remains difficult to engineer into proteins. Here, we explore ligand-induced folding of nanobodies as a general strategy to generate conformational receptors for a broad range of target ligands. More specifically, we have generated partially unfolded nanobody variants that fold exclusively upon binding to their target ligand. To do so, we have introduced destabilizing mutations in the framework region, carefully tuning their stability while preserving molecular recognition. Given the high conservation of the framework across different nanobodies, this approach is broadly applicable to many other target ligands, which has enabled us to engineer conformational switches for three different important biomarkers and drugs: the pregnancy hormone chorionic gonadotropin, the cytokine tumor necrosis factor alpha, and the chemotherapy and immunosuppressant drug methotrexate. Another useful property of conformational signaling is that it can directly gate signal transduction, which we have leveraged to build an optical sensor for the quantitative detection of the pregnancy hormone in urine. Finally, conformational signaling introduces new mechanisms for the allosteric modulation of binding, which we have explored to control the affinity and dynamic range of the sensor. Our work thus establishes a rational strategy for engineering conformational signaling receptors with customizable properties and generalizable for many different target ligands.

**Significance Statement:** Biological receptors often signal by changing conformation upon ligand binding, but this mechanism remains difficult to adapt to compact and generalizable binding scaffolds. Here we show that we can engineer the stability of nanobodies to design conformational receptors, enabling a new class of tunable biosensors and responsive protein devices.

## Introduction

Conformational signaling is a fundamental mechanism by which biological systems sense and respond to molecular cues. In this process, ligand binding induces a structural rearrangement in the receptor, which triggers a functional response. Nature uses this strategy across many different proteins and signaling pathways. Examples include regulating open–closure dynamics of periplasmic binding proteins in bacterial chemotaxis and transport^1^, gating transitions in ligand- and voltage-regulated ion channels^2^, ligand-induced changes in the oligomeric state of receptors and transcription factors^3–5^, and disorder-to-order transitions in intrinsically disordered regions^6–8^.

Beyond its central role in biology, conformational signaling is also useful for biotechnologies, as it can directly convert target binding into a readable output, such as enzyme activation^9^, release of chemicals^10^, change in mechanical properties^11^, or generation of optical or electrochemical responses^12^. And because only binding of the correct ligand at the correct site can satisfy the coordinated network of interactions required to stabilize the signaling conformation, it also provides an additional layer of selectivity, thus helping discriminate true target binding from nonspecific effects in complex biological environments, such as adsorption and background interference^13^. This versatility, selectivity, and specificity make conformational signaling very useful for the design of responsive molecular systems with broad utility in biotechnologies, including biosensors^14,15^, imaging probes^16^, smart biomaterials^17^, and logical actuators^18^.

Engineering conformational signaling into biomolecules remains a challenge in molecular design. Generally, it requires a detailed knowledge of the structure and biophysics of the system, as well as the allosteric networks connecting ligand recognition to a functional output^19,20^. Despite this complexity, several studies have succeeded at controlling already existing intrinsic conformational changes, such as hinge motions in periplasmic binding proteins^21,22^, circular permutation and domain insertion to connect ligand binding to fluorescent or enzymatic readouts^23^, or chemically-induced dimerization systems in which ligand binding drives receptor assembly^24^. Other studies have developed strategies to introduce conformational signaling into otherwise non-conformational receptors, such as domain architectures that shift between competing structural or folding states^25^. Among these, folding-based mechanisms are particularly compelling, as they directly couple molecular recognition to folding, one of the largest conformational changes a protein can undergo. These mechanisms thus offer a powerful strategy for engineering conformational receptors.

Motivated by its potential applications, as well as by its fundamental value for understanding molecular recognition, we set to develop a rational approach to engineer protein-based conformational signaling receptors. To achieve this, we focus on nanobodies, single-domain antibody fragments derived from the ligand-binding domain of the heavy-chain-only antibodies from camelids and sharks^26^. These small, monomeric proteins offer affinity, specificity and versatility comparable to canonical antibodies, but in a mere one tenth of their size. And while nanobodies do not undergo significant conformational changes upon binding to their targets—as neither do antibodies—their modular architecture makes them very convenient for biomolecular engineering. They feature three hypervariable loops responsible for ligand binding, and a highly conserved framework that provides a scaffold responsible for structure and stability. Given this clear separation between the regions responsible for binding and stability, we hypothesize that we can engineer the framework of nanobodies, controlling their biophysical properties without affecting binding affinity and specificity, to convert them into conformational receptors. In particular, given that nanobodies exhibit a two-state—and in many cases reversible—folding^27–29^, we engineer their folding equilibrium to introduce ligand-induced folding. To achieve this, we design marginally unstable nanobody variants, carefully tuning their stability to couple ligand binding to folding, and use this ligand-induced conformational change to generate a measurable signal output that quantitatively informs on ligand concentration.

## Results

### Thermodynamics of Ligand-Induced Folding

Ligand-induced folding couples two chemical equilibria, such that the folding state of the receptor allosterically controls ligand binding (Fig. 1A). Because only the folded state is competent for high-affinity binding, the apparent binding affinity (*K*_D,app_) depends on the intrinsic binding affinity of the folded receptor (*K*_D,int_) as well as on its fold stability (*K*_F_):

**Figure 1.**
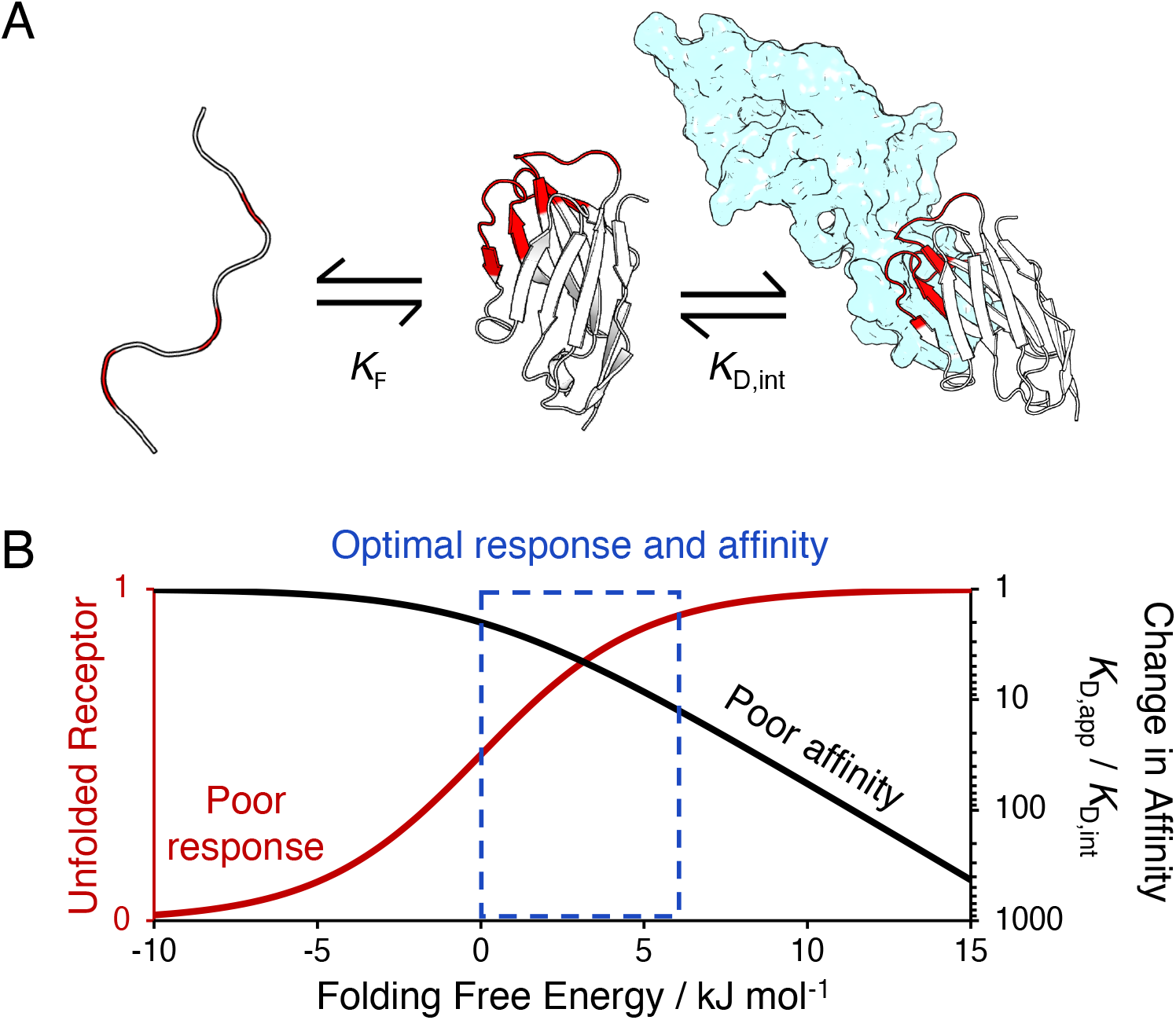
The thermodynamics of ligand-induced folding. (**A**) The ligand-induced folding mechanism couples two chemical equilibria, such that ligand binding (*K*_D,int_) results in folding (*K*_F_), here shown for a nanobody targeting the pregnancy hormone chorionic gonadotropin (structural model of the complex generated with AlphaFold3, nanobody shown as white cartoon with hypervariable loops in red, and hormone in light blue). (**B**) This thermodynamic coupling introduces a trade-off between affinity and signaling, with decreasing protein stability increasing the fraction of unfolded receptor (red line) and thus the population shift upon binding, but at the expense of losing affinity (black line). The optimal compromise lies at the stability range at which there is sufficient unfolded receptor to undergo the ligand-induced folding transition, whereas the penalty in affinity remains low.

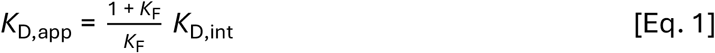

This thermodynamic coupling introduces a trade-oh between affinity and signaling. Destabilizing the receptor increases the unfolded population and thus the magnitude of the ligand-induced population shift, but an excessive destabilization imposes too much of an energetic penalty on ligand binding and thus decreases apparent affinity. Conversely, insufficient destabilization preserves affinity, but reduces the population shift and thus the detectable signal change. Resulting from this, achieving functional ligand-induced folding requires carefully tuning receptor stability to achieve a significant fraction of unfolded molecules, but while retaining favorable binding. In quantitative terms, the optimal trade-off lies at folding free energies in the range of 0–6 kJ mol^−1^, which corresponds to an unfolded population fraction of 50–90% and, resulting from Eq. 1, an affinity within a 2 to 10-fold lower than that of the original stable receptor (Fig. 1B).

### Engineering Nanobody Stability

For the initial optimization, we have focused on an already existing nanobody against human chorionic gonadotropin (hCG), a female reproductive hormone secreted by the placenta. This 50 kDa glycoprotein is widely used a clinical biomarker of pregnancy in commercial pregnancy tests. These tests provide a largely qualitative, yes/no response, which is often enough to detect pregnancy. However, hCG is also a biomarker of embryo implantation during pregnancy, gestational trophoblastic diseases, as well as ovarian and testicular tumors^30^. The development of sensors capable of quantitative and dynamic hCG monitoring will thus expand its clinical utility, enabling for example at-home monitoring of placental development and embryo implantation in high-risk pregnancies. To that end, conformational signaling offers a convenient sensing strategy, because it directly couples molecular recognition to signal transduction in a rapid, reversible, and potentially reagent-less format.

We set up to engineer the anti-hCG nanobody into a ligand-induced folding conformational switch. Aiming for a controlled nanobody destabilization while preventing binding disruption, we introduced destabilizing mutations in the conserved framework region, preserving the antigen-binding hypervariable loops. To achieve this, we have pursued two alternative strategies: (i) removing a conserved disulfide bond by mutating cysteines 22 and 92 to Ala, Gly, Ser, Met and Trp, and (ii) introducing cavity-creating mutations to weaken the hydrophobic core of the protein by substituting Val, Ile, Leu and Trp residues with Ala (Fig. 2A). Removing the disulfide bond decreases stability by increasing the conformational entropy of the unfolded state^31^, whereas cavity-creating amino acid substitutions replace a bulky side chain with a smaller one, generating an internal cavity whose volume correlates with the extent of thermodynamic destabilization through loss of packing^32^. We predicted the stability of engineered variants generated using these two strategies with a computational pipeline combining structure prediction with AlphaFold2 and energetic scoring with Rosetta^33^. The predicted changes in thermodynamic stability for 13 single and double mutants of the anti-hCG nanobody (8 removing the disulfide bond and 5 creating a cavity at the hydrophobic core) are shown in Fig. S1. While no clear trend in the predicted thermodynamic stabilities is observed for the disulfide variants, the cavity-creating mutations show a dependence of predicted stability on the size of the bulky side chain being replaced, with the largest destabilization corresponding to mutation of the bulkier Trp37, in line with the expected trend of increased destabilization with increasing cavity size^32^.

**Figure 2.**
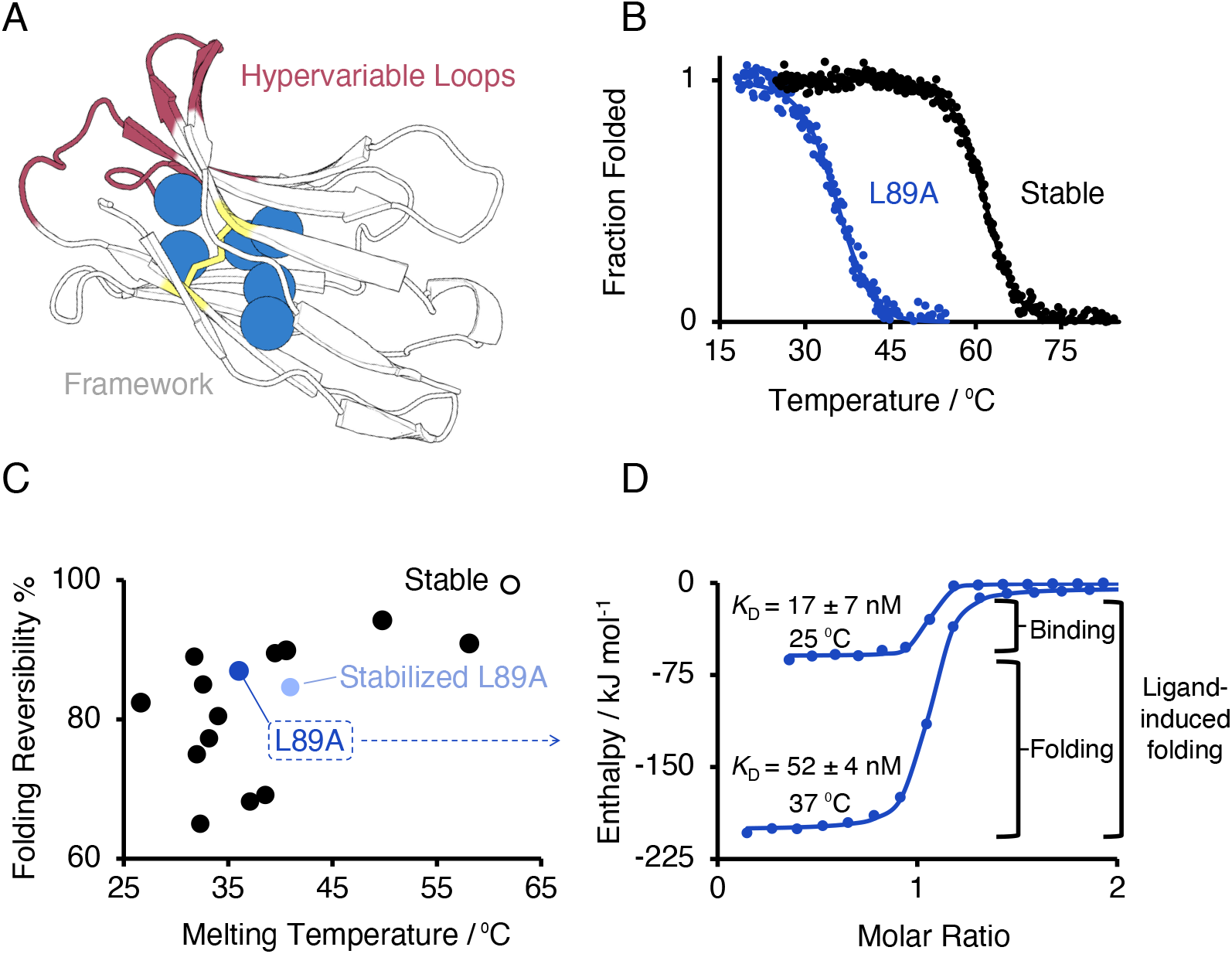
Engineering ligand-induced folding into the anti-hCG nanobody. (**A**) We destabilize the framework region (white cartoon) either by introducing cavity-creating mutations (blue spheres) or by eliminating the conserved disulfide bond (yellow sticks), preserving the hypervariable loops (in red) to minimize binding disruption. (**B**) We then assessed the folding stability and reversibility of these variants by thermal denaturation experiments monitored by circular dichroism. (**C**) We tested the L89A variant as an initial candidate for ligand-induced folding, with a melting temperature of 35.7ºC and a folding reversibility of 87%. (**D**) Comparison of isothermal titration calorimetry experiments for this variant at 37 ºC (a temperature at which it is 60% unfolded) and 25 ºC (completely folded) shows nanomolar affinity for hCG, supporting our hypothesis that framework engineering enables controlling stability without significantly affecting binding. We also observe a much higher exothermic release, consistent with the coupling of binding to folding.

We produced these 13 destabilized variants, together with the original stable nanobody for experimental validation, evaluating their stability and folding reversibility by thermal denaturation experiments monitored by circular dichroism (Fig. 2B; Table S1). Our goal is to identify nanobody variants suitable for ligand-induced folding. More specifically, we aim for a thermal denaturation midpoint slightly below 37 ºC—for achieving functional conformational switching at body temperature—and good folding reversibility. We observed a broad range of stabilities, with melting temperatures ranging from 61 ºC for the original nanobody to 27 ºC for the most destabilized variant (Fig. 2C). For the cavity-creating variants, we find a good correlation between the experimental values and computational predictions. For the disulfide variants, in contrast, the correlation is significantly worse (Fig. S1). This poorer correlation is likely due to the fact that removal of the disulfide bond affects the conformational space of the unfolded state, which is not modeled explicitly but is instead accounted for through a per-residue additive term in the Rosetta energy function^34^. We have thus focused on the cavity-creating approach, as it provides a more predictable engineering strategy and a wider range of stabilities. We thus selected the variant bearing the L89A mutation. Truncation of this core leucine into alanine decreases the thermal denaturation midpoint by 26 ºC to 35.7 ± 0.1 ºC, resulting in the unfolding of 60% of the nanobody at 37 ºC—right at the optimal range for ligand-induced folding—while retaining a remarkable 87% folding reversibility (Fig. 2C). Consistent with this, NMR experiments showed partial nanobody unfolding at 37 ºC (Fig. S2).

### Evaluation of Ligand-Induced Folding

We next evaluated the binding of the engineered L89A nanobody variant by performing isothermal titration calorimetry (ITC) experiments. First, we evaluated binding at 25 ºC, a temperature at which the nanobody is fully folded, observing a dissociation constant (*K*_D,int_) of 17 ± 7 nM, within error of the 13 ± 9 nM of the original nanobody (Fig. S3). Upon increasing the temperature to 37 ºC the affinity of the original nanobody remains unchanged. In contrast, the affinity of the engineered nanobody decreases by a 3-fold to 52 ± 4 nM, very close to the 2.5-fold affinity reduction predicted by the coupling of folding and binding (Fig. 1; Eq. 1). Notably, the engineered nanobody features a much higher exothermic signature. More specifically, the excess 140 kJ mol^−1^ enthalpy we observe for this L89A variant is consistent with our expected estimate of 107 kJ mol^−1^ for the folding of a 60% unfolded nanobody (details on the calculation in supplementary information and Fig. S4). Also consistent with these calorimetric results, NMR experiments are also consistent with nanobody folding upon binding to hCG (Fig. S2). These results thus validate our design hypothesis of engineering the nanobody framework to control its biophysical properties without significantly affecting molecular recognition, as well as our mechanistic model of engineering framework stability to couple binding with folding.

### Generalization to Other Nanobodies

Given the high conservation of the framework region across different nanobodies, we hypothesize that our approach can be generalizable to produce conformational switching receptors for other ligands. To demonstrate this, we applied the same approach to two other nanobodies targeting tumor necrosis factor alpha (TNFa), a proinflammatory cytokine used as a biomarker of immune response, inflammation, and infection^35^; and methotrexate (MTX), an immunosuppressant and chemotherapy drug requiring therapeutic drug monitoring^36^. We introduced some of the mutations identified for the anti-hCG nanobody and measured their effect on stability. We find that, although the absolute stability depends on the intrinsic stability of each nanobody, the relative order of the destabilizing effect of the mutations remains consistent across the three nanobodies (Table S2). We then selected variants showing appropriate destabilization and folding reversibility to evaluate their binding (Fig. S5). Again, we find a larger exothermic fingerprint, consistent with the coupling of binding and folding, and a dependence of binding affinity with folding stability, consistent with the ligand-induced folding thermodynamic model (Fig. 3). These results thus support the general transferability of the approach across different nanobodies targeting other ligands.

**Figure 3.**
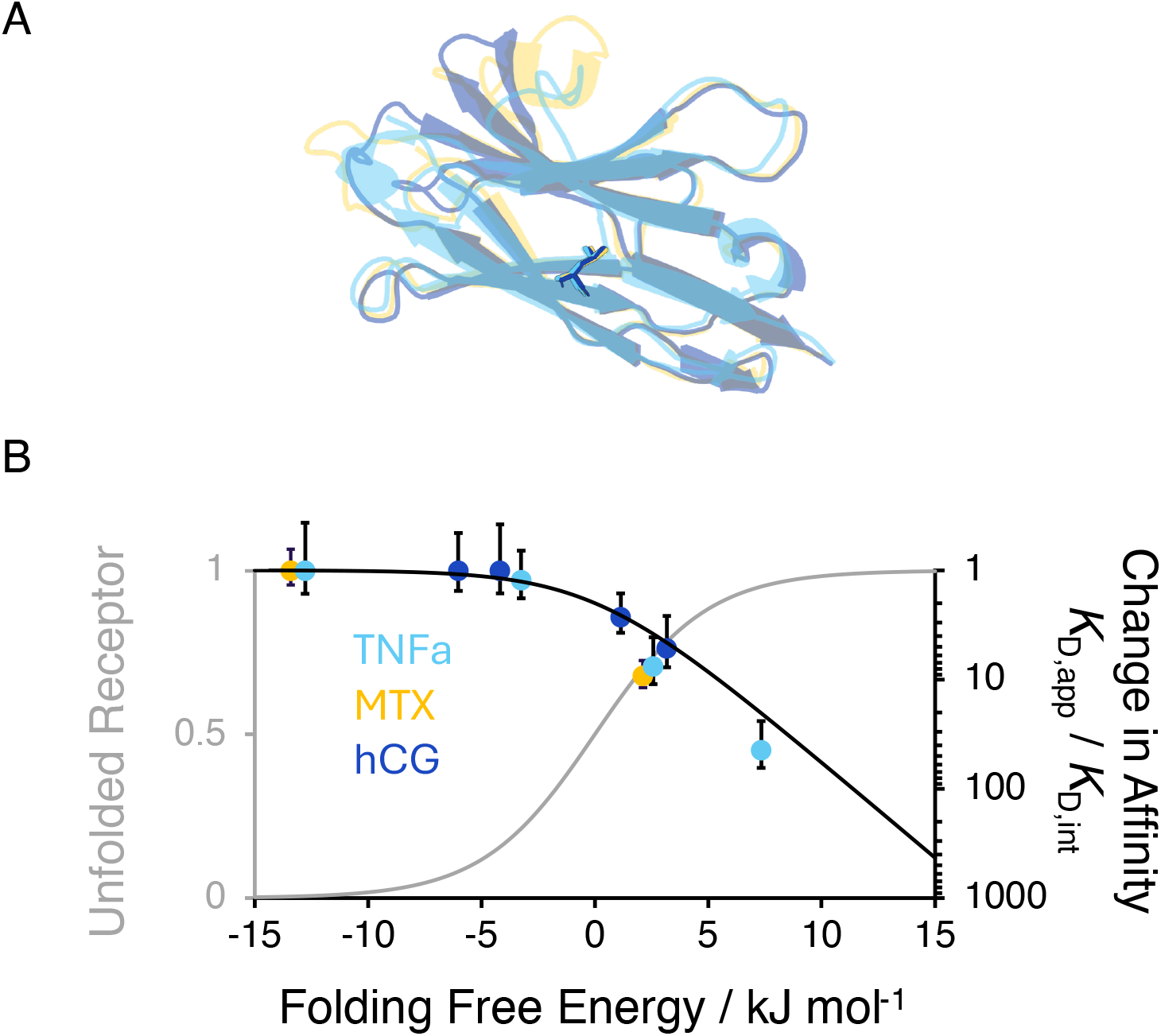
Transferability of conformational signaling across different nanobodies. (**A**) We have transferred some of the destabilizing mutations (leucine 89 highlighted in sticks as an example of the high sequence and structural conservation) of the anti-hCG scaffold (dark blue) to two other nanobodies targeting the cytokine tumor necrosis factor alpha (TNFa, light blue) and the drug methotrexate (MTX, yellow). (**B**) Similar to the anti-hCG nanobody, these variants support ligand-induced folding, with their affinity following the expected dependence on folding stability (Eq. 1; Fig. 1B).

### Implementation in an Optical Biosensor for Detection of hCG in Urine

To explore whether conformational signaling can directly gate signal transduction, we developed an optical readout based on the anti-hCG nanobody L89A conformational receptor. Specifically, we substituted Ser7 by a cysteine—a position distant from the antigen-binding region yet close in sequence to tryptophans—and modified its free thiol with a maleimide-Alexa488 fluorophore. In the absence of hCG, the exposed aromatic side chains of the unfolded nanobody quench the fluorescence. In the presence of hCG, in contrast, folding of the nanobody buries the aromatic residues and activates fluorescent emission^37^. First, we tested the sensor in phosphate buffer at 37 ºC. Titration with hCG produced an increase in fluorescence proportional to hCG concentration, following a hyperbolic dependence that we can fit to a Langmuir binding isotherm with an apparent dissociation constant of 100 ± 30 nM (Fig. S6). In contrast, when we performed the same experiment at 25 ºC, a temperature at which this variant is fully folded, we did not observe a significant change in fluorescence, confirming that the signaling strictly arises due to the coupling of binding with the folding (Fig. S6). We next evaluated sensor performance in real human urine, a more relevant matrix for hCG testing. Again, we observe a clear fluorescence increase proportional to hCG concentration, with an apparent dissociation constant of 40 ± 5 nM (Fig. 4B)

**Figure 4.**
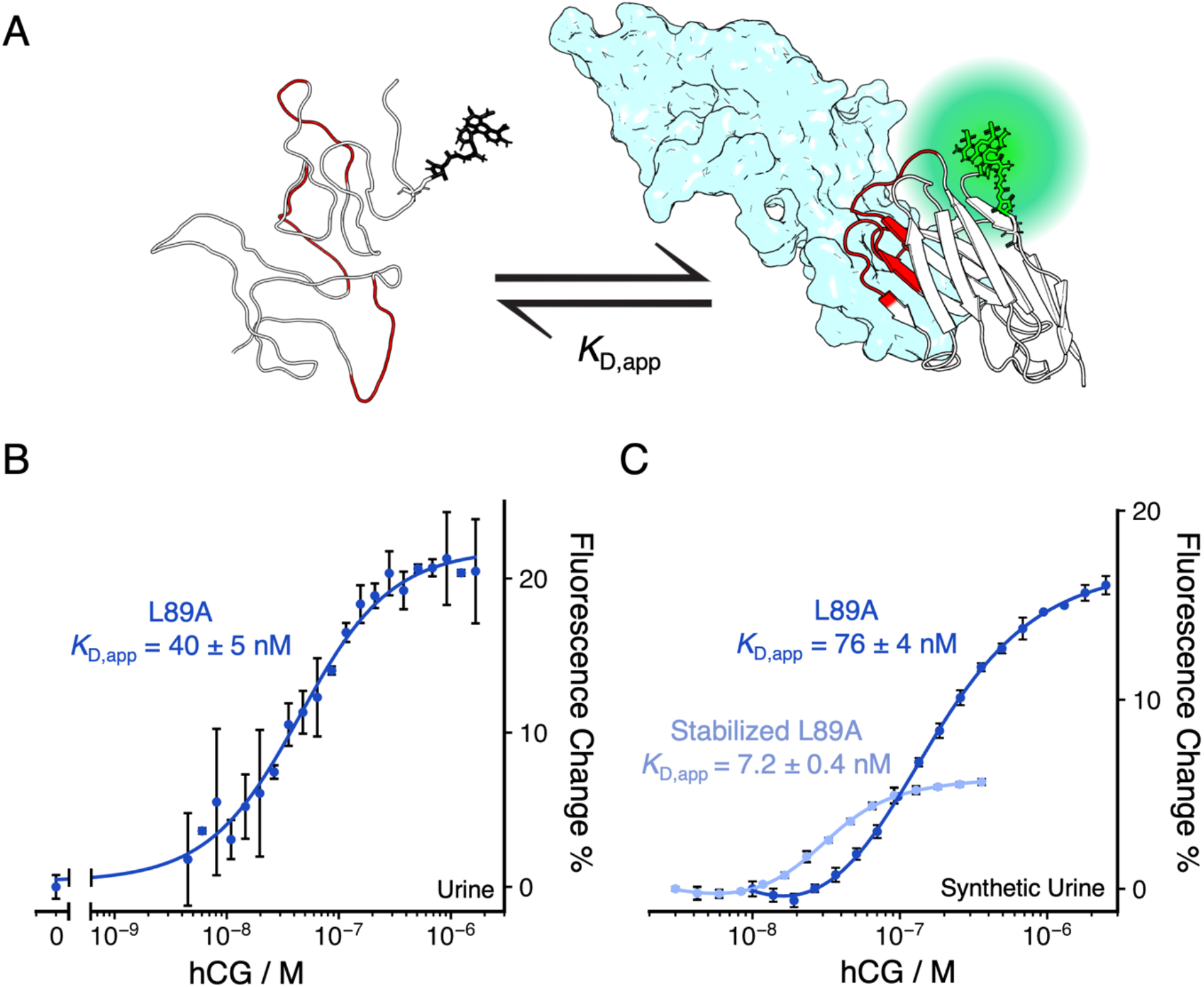
Optical sensor for measuring pregnancy hormone in urine. (**A**) We labelled the anti-hCG L89A nanobody with an Alexa-488 fluorescent reporter (represented as green sticks) at a position sensitive to the folding of the nanobody. Nanobody unfolding in the absence of hCG quenches fluorescence, whereas folding upon hCG binding activates emission. (**B**) We tested the sensor in real human urine at 37 °C by titrating increasing concentrations of hCG. In doing this, we observed an increase in fluorescent signal and a dissociation constant of 40 ± 5 nM (circles represent experimental measurements, and solid line the best fit to a Langmuir isotherm model) (**C**) We can allosterically modulate the affinity of the sensor by using variants with different stabilities. For example, a variant incorporating the L89A mutation into a stabilized nanobody scaffold (light blue) shows better affinity than the L89A variant (dark blue), but lower signal change, as expected due to its higher stability. Error bars represent the confidence intervals for three technical replicates.

### Allosteric Control of Binding through Stability

The coupling of binding and folding implies that we can tune the conformational signaling of the sensor by modulating nanobody folding equilibrium. We thus reason that by using variants with different stabilities we can develop biosensors spanning across different dynamic ranges. To test this idea, we built a sensor using instead a variant incorporating the same L89A mutation on a nanobody scaffold previously stabilized with ProteinMPNN^38^. With a melting temperature of 41 °C, this variant is slightly more stable than the L89A variant used in the previous experiments, showing as well similarly high unfolding reversibility (Fig. 2C). We functionalized it with the same fluorophore labeling strategy already described and titrated it with hCG at 37 ºC (Fig. 4C). Again, we observe an increase in fluorescent signal dependent on hCG concentration. We observe, however, an improvement in affinity with respect to the L89A variant, with a 10-fold lower dissociation constant of 7.2 ± 0.4 nM. The additional stability of this variant can thus bring the detection window to concentrations in the low nanomolar range, while still being sufficiently destabilized to produce a signaling output. However, as expected from the affinity–signaling trade-oh, the magnitude of the signal change is nearly one third of the L89A variant alone. These results thus highlight the potential of ligand-induced folding mechanism to allosterically control sensor’s affinity and dynamic range through the rational tuning of folding stability.

## Discussion

In this study, we have developed molecular switches based on nanobodies, exploring a general conformational signaling strategy based on the thermodynamic coupling of binding and folding. Researchers have used folding-based strategies to engineer structure-switching nucleic acids and proteins such as, for example, ligand-induced folding^37^, mutually exclusive folding^25^, or alternate-frame folding^39^. In all these cases, functional signaling depends on an energetic balance, by which destabilization of the binding-competent state enhances the ligand-driven population shift and thus signaling effciency. This mechanism, however, unavoidably incurs in an energetic penalty due to the cost of coupling recognition to folding, introducing an inherent trade-off between affinity and response. And while this may render the approach less suitable to some specific applications demanding very low detection limits, in practice the penalty remains reasonable. For example, as long as the unfolded receptor population remains within the 50-90% range—equivalent to a stability of 0-6 kJ mol^−1^—the affinity decreases by less than one order of magnitude. Consistent with this, our engineered receptors achieve affinities within 3–50-fold lower than those of the original stable nanobodies.

This thermodynamic coupling between molecular recognition and folding introduces a mechanism for allosteric control^40^. The affinity of the conformational switch reflects both the binding free energy of the fully folded receptor as well as the folding free energy necessary to populate the binding-competent state. We can thus customize the affinity of the receptors by engineering their stability. This flexibility can be useful, for example, to match a specific detection threshold, or to adapt to complex sample matrices such as urine or other biofluids. It can also support multiplexing, combining receptors with different affinities to expand the analytical detection range^41^. Typical affinity receptors exhibit a 81-fold dynamic range—as dictated by the Langmuir isotherm^40^—which limits their use in applications demanding broader dynamic ranges. The levels of hCG, for example, vary by as much as four orders of magnitude during the first trimester of pregnancy. Pregnancy tests, however, typically use very high affinity antibodies to lower the detection limit of hCG, aiming to detect pregnancy as early as possible. But in exchange, their response saturates shortly after the first four weeks. In contrast, biosensors with a customizable dynamic range would cover a wider range of concentrations, enabling hCG monitoring throughout the entire pregnancy.

Nanobodies are particularly suitable for engineering ligand-induced folding because of their compact format, modular architecture, and reversible folding. This facilitates engineering their biophysical properties without affecting affinity and specificity. In prior work, we and others have leveraged this idea by redesigning the framework of nanobodies to improve their thermostability and production^38,42,43^. Here, in contrast, we have successfully identified a diverse set of destabilizing mutations to generate conformational switches, enabling systematic and rational control of the conformational equilibrium. We have found, however, that not all mutations allow this. For example, while the variants targeting the disulfide bond achieve sufficient destabilization and reversible thermal unfolding^44^, they generally are less suitable for ligand-induced folding, with only one out of the 11 disulfide variants tested yielding reversible ligand-induced folding (anti-MTX MPNN C22S/C92S variant, table S2). This suggests that the pathway followed upon thermal unfolding differs from ligand-induced folding, with the latter being more reliant on the conserved disulfide bond. In contrast, mutations trimming the hydrophobic core enable a more predictable and rational destabilization.

The strategy presented here complement other approaches to engineer ligand-induced folding. Prior work from the Plaxco lab, for example, introduced terminal truncations on the Fyn SH3 tyrosine kinase domain to couple folding to the binding of its target peptide^37^. Terminal truncations, however, significantly compromise protein expression and solubility. Later work by the Koder lab successfully leveraged surface electrostatic repulsion to engineer supercharged scFv antibody fragments supporting ligand-induced folding^45^. This approach can be more easily generalized to scFv targeting other antigens and, potentially, to other proteins^46^. However, it requires a significantly higher number of substitutions (about 10 % of amino acids) and introduces a strong dependence of stability on ionic strength. Finally, work by the Cepko lab developed conditionally stable nanobodies^47^, but their reliance on vast variant libraries requires complex large-scale functional screenings, and do not provide thermodynamic information to rationally control the conformational switching equilibrium. Our engineering approach, in contrast, preserves protein expression and solubility, relies on well-established thermodynamic principles enabling rational and predictable design, and is simpler to implement, often requiring just a single substitution. In addition, given the high sequence and structural conservation across nanobody frameworks, our design rationale and substitutions are generally transferable to different targets, such as the anti-hCG, anti-TNFa and anti-MTX nanobodies tested here. We also hypothesize that the same ligand-induced folding mechanism may apply to other families of single-domain, monomeric, and engineerable receptors, including for example adnectins^48^, affibodies^49^, or even artificial proteins.

Finally, a useful property of conformational signaling is that it can provide straightforward signal transduction. If the conformational change is large enough, such as folding/unfolding, it can directly gate transduction of binding into a measurable output signal. This can facilitate the development of new biotechnological applications. Here, we have built an optical sensor for the pregnancy hormone. But we hypothesize that this mechanism can also support expansion to other signal transduction strategies, such as electrochemical, enzymatic, mechanical, plasmonic, or transcriptional. The combination of generalizable receptors, signaling mechanisms, and design strategies offers a powerful approach towards the design of conformational switches, potentially useful in applications in diagnosis, therapy, and research, such as continuous monitoring biosensors, chemical-responsive smart biomaterials, and biosynthetic circuits.

## Conclusion

In this work, we have engineered nanobodies into conformational receptors using ligand-induced folding. By introducing destabilizing mutations into the framework region, we have achieved a rational and controlled destabilization, coupling binding with folding without affecting ligand recognition. We have then implemented a direct optical signal transduction, building a sensor for the detection of the pregnancy hormone hCG in urine. Furthermore, we have shown that by tuning the stability of the nanobody we can allosterically control the affinity and dynamic range of the sensor. We have also demonstrated the general applicability of this approach, by designing conformational switches based on two other unrelated nanobodies against the cytokine TNFa and the drug methotrexate. In conclusion, our results establish a practical and transferable strategy for engineering ligand-responsive nanobody switches, with potential application in the development of biosensors and other biotechnologies.

## Acknowledgements

This work was supported by the Spanish Agencia Estatal de Investigación and the Spanish Ministry of Science and European Union NextGenerationEU/PRTR (grants PID2023-148199OA-I00, PID2024-160774OB-I00, PID2025-174578NA-I00, RYC2021-031074-I, RYC2022-036457-I). GOQ, FP, and GJ-O acknowledge financial support by Ikerbasque, the Basque Foundation for Science. GJ-O declares he is a founder and chief scientific officer of Protellix Biotech, which is interested in developing nanobodies for diverse biotechnological applications. The rest of the authors declare no other competing interests.

