## Supplementary material for "Engineering Nanobody Conformational Switches": Materials and Methods

### Materials and Methods for:

**Computational prediction of nanobody stability:**

We predict the change in thermodynamic stability of the anti-hCG nanobody following five single cavity-creating mutations (L21A, W37A, I71A, V80A, L82A) and single and double mutations targeting the disulfide bond between C22 and C92 (C22A, C22G, C22A/C92A, C22A/C92M, C22G/C92A, C22G/C92M, C22G/C92W, and C22S/C92S) using a computational pipeline based on Rosetta energies averaged over AlphaFold2 ensembles and validated on several enzyme families[1].

In short, for each mutant and the original nanobodies, we use the AlphaFold 2.3.2 version[2] to generate 100 structures using model 3 and the following options:

```
--db-preset=reduced_dbs
--model-preset=monomer
--max-template-date 2050-01-01
--enable-gpu-relax
```

Then, each structure is relaxed with the *minimize* application of Rosetta version 3.4.0[3] with the following flags:

```
-run:min_type lbfgs_armijo_nonmonotone
-run:min_tolerance 0.001
```

The folding free energy $\Delta G_f$ of each variant and the wild type is computed as the average of its 25 lowest-energy values. The $\Delta\Delta G_f$ value for each variant is then computed as the difference between the $\Delta G_f$ of the variant and that of the wild type, with positive and negative $\Delta\Delta G_f$ values indicating diminished and increased stability, respectively.

**Protein expression and purification in bacteria:**

We introduce the gene sequences for the nanobodies into an ampicillin-resistant pET3a plasmid between the NdeI and BamHI restriction sites, ordering the gene synthesis and cloning from commercial sources (GenScript). We then transform the plasmids in BL21(DE3) Escherichia coli (New England BioLabs) and grow them in LB media (Condalab) at 37 °C until an OD600 of 0.6, induce protein overexpression by adding 1 mM IPTG, and incubate at 20 °C overnight. When expressed in bacteria, nanobodies can form inclusion bodies. To recover the fraction of nanobodies in inclusion bodies, we perform a chemical refolding protocol. Specifically, we resuspend the pellet in buffer with 6 M guanidinium chloride (Calbiochem-Omnipur) and dialyze overnight against native buffer at 4 ºC. We then separate the supernatant by centrifugation and apply it to a Ni-NTA resin (Millipore) for affinity chromatography purification, wash it with 25 mM imidazole, and elute with 250 mM imidazole. Finally, we purify the eluate by size exclusion chromatography on a HiLoad 26/600 Superdex 75 pg column (Cytiva). We assess protein purity by SDS-PAGE, and measure absorbance at 280 nm to determine the total amount of protein obtained.

**Thermostability measurements:**

We perform thermal denaturations monitored by circular dichroism (CD) on a J-850 spectrophotometer (JASCO). To do this, we prepare a 5-15 μM protein solution in native buffer in a 2 mm cuvette. We then increase the temperature at 1 degree per minute, recording in 0.1 degree intervals the ellipticity at 210-222 nm with a bandwidth of 4 nm and digital integration time of 8 sec. For each nanobody we perform consecutive heating and cooling experiments, using the changes in ellipticity to monitor the protein unfolding and refolding equilibrium. We then analyze the experimental data using a non-linear least-squares minimization method to fit the experimental ellipticities to the Gibbs-Helmholtz equation for a two-states reversible folding equilibrium:

$$\theta = \frac{\theta_{\mathrm{F}} + m_{\mathrm{F}}\,\mathrm{T} + (\,\theta_{\mathrm{U}} + m_{\mathrm{U}}\,\mathrm{T}\,)\,\mathrm{e}^{-\Delta G/_{\mathrm{RT}}}}{1+\mathrm{e}^{-\Delta G/_{\mathrm{RT}}}} \quad \text{Eq. 1}$$

$$\Delta G = \Delta H_{\mathrm{m}}\left(\frac{T_{\mathrm{m}} - \mathrm{T}}{T_{\mathrm{m}}}\right) + \Delta C_{\mathrm{P}}\left(\mathrm{T} - T_{\mathrm{m}} - \mathrm{T}\,\mathrm{Ln}\frac{\mathrm{T}}{T_{\mathrm{m}}}\right) \quad \text{Eq. 2}$$

We extract six parameters: the intrinsic ellipticities in the absence of denaturant for the folded and unfolded states, $\theta_{\mathrm{F}}$ and $\theta_{\mathrm{U}}$; the slopes for the linear dependence of the ellipticities on temperature for the folded and unfolded states, $m_{\mathrm{F}}$ and $m_{\mathrm{U}}$; the melting temperature, $T_{\mathrm{m}}$; and the enthalpy of melting, $\Delta H_{\mathrm{m}}$. We calculate the change in heat capacity, $\Delta C_{\mathrm{P}}$, using the empirical correlations reported by Myers et al[4].

**Isothermal titration calorimetry:**

We evaluate anti-hCG, anti-MTX and anti-TNFa nanobody binding affinity by using a MicroCal PEAQ-ITC (Malvern Panalytical). We purchased hCG hormone (MP Biomedicals, Ref. 02198591-CF) and methotrexate hydrate (Sigma-Aldrich, Ref. A6770) from commercial sources. For TNFa, we purchased a pET-11d plasmid encoding recombinant His-tagged TNFa (GenScript) for production in E. coli, purify it by Ni-NTA affinity chromatography and size exclusion, and verify its purity by SDS-PAGE. We fill the cell with a buffered solution of 5–30 μM nanobody and titrate it with a buffered solution of 50–300 μM of the corresponding molecular target. For the anti-hCG nanobody, due to the scarcity of the hormone, we invert the setup, filling the cell with 20 μM hCG and the syringe with 190 μM nanobody. We perform consecutive target additions, waiting a minimum of 100 second interval between additions for complete equilibration, until saturation. We integrate and analyze the corresponding calorimetry curves using the MicroCal PEAQ-ITC Analysis Software, fitting the data to a single-site binding isotherm to obtain values for the dissociation constant and binding enthalpy. Reported uncertainties correspond to 95% confidence intervals derived from the fit.

**Fluorescence measurements:**

We design and produce variants of the hCG nanobody incorporating a cysteine (S7C), which we conjugate to a maleimide-modified fluorophore ATTO488 (ATTO-TEC) via a Michael addition. Specifically, we mix a solution of 40-60 μM of nanobody with a 5-fold excess of dye in 50 mM sodium phosphate buffer at pH 7.0, with 100 mM NaCl, and incubate at a room temperature for 3 hours.

To perform the fluorescence measurements, we use a JASCO-6500 spectrofluorometer and a 10 mm x 4 mm quartz cuvette (Hellma Analytics). We measure the fluorescence signal using an excitation wavelength of 504 nm and recording the emission signal between 513 and 530 nm with a 3 nm bandwidth. We performed titrations using logarithmic equidistant hormone concentrations and waiting 2 minutes for equilibration under continuous agitation. For each titration point, we calculate the fluorescence intensity as the average signal within the selected wavelength interval surrounding the emission maximum (518–520 nm) from a total of three spectrum accumulations.

We analyzed the data using custom Python scripts, fitting the data to a Langmuir isotherm by nonlinear least-squares minimization to extract the affinity. More specifically, we describe fluorescence ($F_{obs}$) as the population-weighted addition of both fluorescence of the free ($F_{free}$) and bound ($F_{bound}$) states as:

$$F_{\mathrm{obs}} = F_{\mathrm{free}}(1 - f_{\mathrm{b}}) + F_{\mathrm{bound}}\, f_{\mathrm{b}} \qquad \text{Eq. 3}$$

Where the population fraction of bound nanobody ($f_b$) corresponds to:

$$f_{\mathrm{b}} = \frac{[\mathrm{P}]_{\mathrm{T}} + [\mathrm{L}] + K_{\mathrm{D,app}} - \sqrt{\left([\mathrm{P}]_{\mathrm{T}} + [\mathrm{L}] + K_{\mathrm{D,app}}\right)^2 - 4\,[\mathrm{P}]_{\mathrm{T}}\,[\mathrm{L}]}}{2\,[\mathrm{P}]_{\mathrm{T}}} \qquad \text{Eq. 4}$$

Where $K_{\mathrm{D,app}}$ is the apparent dissociation constant, $[\mathrm{P}]_{\mathrm{T}}$ is the total nanobody concentration in the cell, and $[\mathrm{L}]$ the added concentration of hCG at each titration point.

We also account for the linear dependence of the intrinsic fluorescence of free and bound states on ligand concentration:

$$F_{\mathrm{free}} = b_{\mathrm{free}} + m_{\mathrm{free}} \log{[\mathrm{L}]} \;\; ; \qquad F_{\mathrm{bound}} = b_{\mathrm{bound}} + m_{\mathrm{bound}} \log{[\mathrm{L}]} \qquad \text{Eq. 5}$$

Where $b_{free}$ and $b_{bound}$ are the fluorescence intercepts, and $m_{free}$ and $m_{bound}$ the slopes, for the ligand-free and ligand-bound states, respectively.

**NMR measurements:** We prepared 300 μL samples containing 83 μM $^{15}N$-labelled anti-hCG nanobody L89A variant and 8% (v/v) $D_2O$ in Shigemi microtubes (Shigemi Co., Ltd., Tokyo, Japan). Two-dimensional $^{1}H$–$^{15}N$ heteronuclear single-quantum coherence (HSQC) spectra were acquired on a Bruker Avance III 800 MHz spectrometer equipped with a TCI cryoprobe. We first recorded a spectrum of the free nanobody at 25 °C to characterize its folded state. We subsequently increased the temperature to 37 °C and acquired a second HSQC spectrum under identical conditions to evaluate the temperature-induced change in the conformational state. To evaluate antigen-induced folding, we prepared a separate NMR sample containing $^{15}$N-labelled anti-hCG nanobody, incubate it at 37 ºC for 5 minutes, and add hCG (also pre-heated at 37 ºC) to a 1:1 molar ratio. We acquired a $^{1}$H-$^{15}$N HSQC spectrum of the nanobody–hCG complex at 37 °C under the same experimental conditions used for the free nanobody.

**Chemical denaturation measurements:**

To calculate protein unfolding free energies, we performed chemical denaturations using guanidinium chloride, which we monitored by electronic circular dichroism (ECD) using a J-850 spectrophotometer (JASCO). We prepared a 2 μM protein solution in 50 mM sodium phosphate buffer at pH 6.0. We then titrated this solution with guanidinium chloride in a quartz cuvette of 1 cm path length.  After each guanidinium chloride addition, we stirred the sample for 1 min before measurement to allow for sufficient equilibration. We recorded circular dichroism spectra from 218 to 300 nm at 100 nm/min with a bandwidth of 4 nm, digital integration time of 4 s, and accumulating 4 scans. We used changes in ellipticity at 228–238 nm to monitor the protein unfolding equilibrium.

We analyzed the data using custom Matlab scripts to fit the denaturation curves to a two-state folding equilibrium model following the linear extrapolation method (Santoro & Bolen, 1988). We used non-linear least-squares minimization to fit the circular dichroism ellipticities to the following equations:

$$I_{\mathrm{obs}} = \frac{I_{\mathrm{F}} + I_{\mathrm{U}}\,\mathrm{e}^{\frac{-\Delta G^{\circ}{}_{\mathrm{U}}(0) - m_{\mathrm{G}}\,[\mathrm{GuHCl}]}{\mathrm{RT}}}}{1 + \mathrm{e}^{\frac{-\Delta G^{\circ}{}_{\mathrm{U}}(0) - m_{\mathrm{G}}\,[\mathrm{GuHCl}]}{\mathrm{RT}}}} \qquad \text{Eq. 6}$$

$$I_{\mathrm{F}} = I_{\mathrm{F}}(0) + m_{\mathrm{F}}[\mathrm{GuHCl}]; \qquad I_{\mathrm{U}} = I_{\mathrm{U}}(0) + m_{\mathrm{U}}[\mathrm{GuHCl}] \qquad \text{Eq. 7}$$

We extract six parameters: the intrinsic ellipticity of the folded state in the absence of denaturant, $I_{\mathrm{F}}(0)$; the intrinsic ellipticity of the unfolded state in the absence of denaturant, $I_{\mathrm{U}}(0)$; the slope for the linear dependence of the ellipticity of the folded state on guanidinium chloride concentration, $m_{\mathrm{F}}$ ; the slope for the linear dependence of the ellipticity of the unfolded state on guanidinium chloride concentration, $m_{\mathrm{U}}$; the unfolding free energy in the absence of denaturant, $\Delta G^{\mathrm{o}}{}_{\mathrm{U}}(0)$; and the slope of the linear dependence of the unfolding free energy with the denaturant concentration, $m_{\mathrm{G}}$.

We measured the temperature dependence of the unfolding free energy for the parent nanobodies by performing guanidinium chloride chemical denaturations at different temperatures. We then reconstruct the temperature-stability dependence by fitting this data, along with the stabilities calculated from the thermal denaturation data in the range of 56–67 ºC, to the Gibbs–Helmholtz equation (Eq. 2). From this analysis, we extract the melting temperature, $T_{\mathrm{m}}$; the unfolding enthalpy, $\Delta H_{\mathrm{m}}$; and the change in heat capacity upon unfolding, $\Delta C_{\mathrm{P}}$ (see supplementary information for additional details).
