## Supplementary Information for "Engineering Nanobody Conformational Switches"

**Contents:**

Figures S1 to S6

Tables S1 to S3

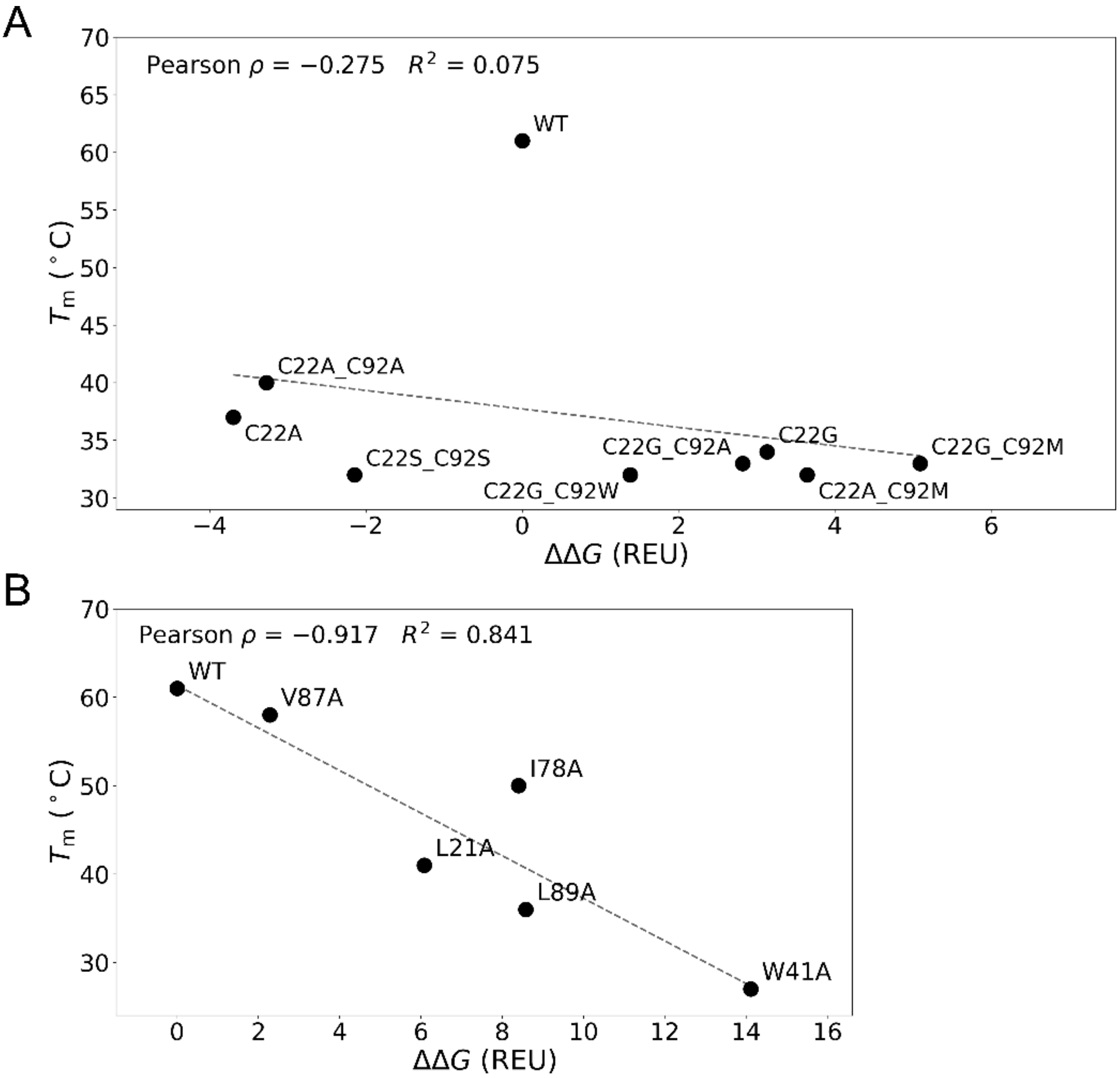


**Figure S1. Predicted *versus* experimental thermal stability of anti-hCG nanobody variants.** (**A**) Single and double mutations removing the disulfide bond between cysteines 22 and 92. (**B**) Cavity-creating mutations. In both cases, the predicted thermodynamic (de)stabilization is expressed as the $\Delta\Delta G_f$ relative to the wild type in Rosetta energy units (REU), and the thermal stability as the melting temperature ($T_m$) determined experimentally, with Pearson's correlation coefficient (ρ) and the coefficient of determination ($R^2$) indicated. The dashed lines represent least-squares linear fits to all plotted data points.

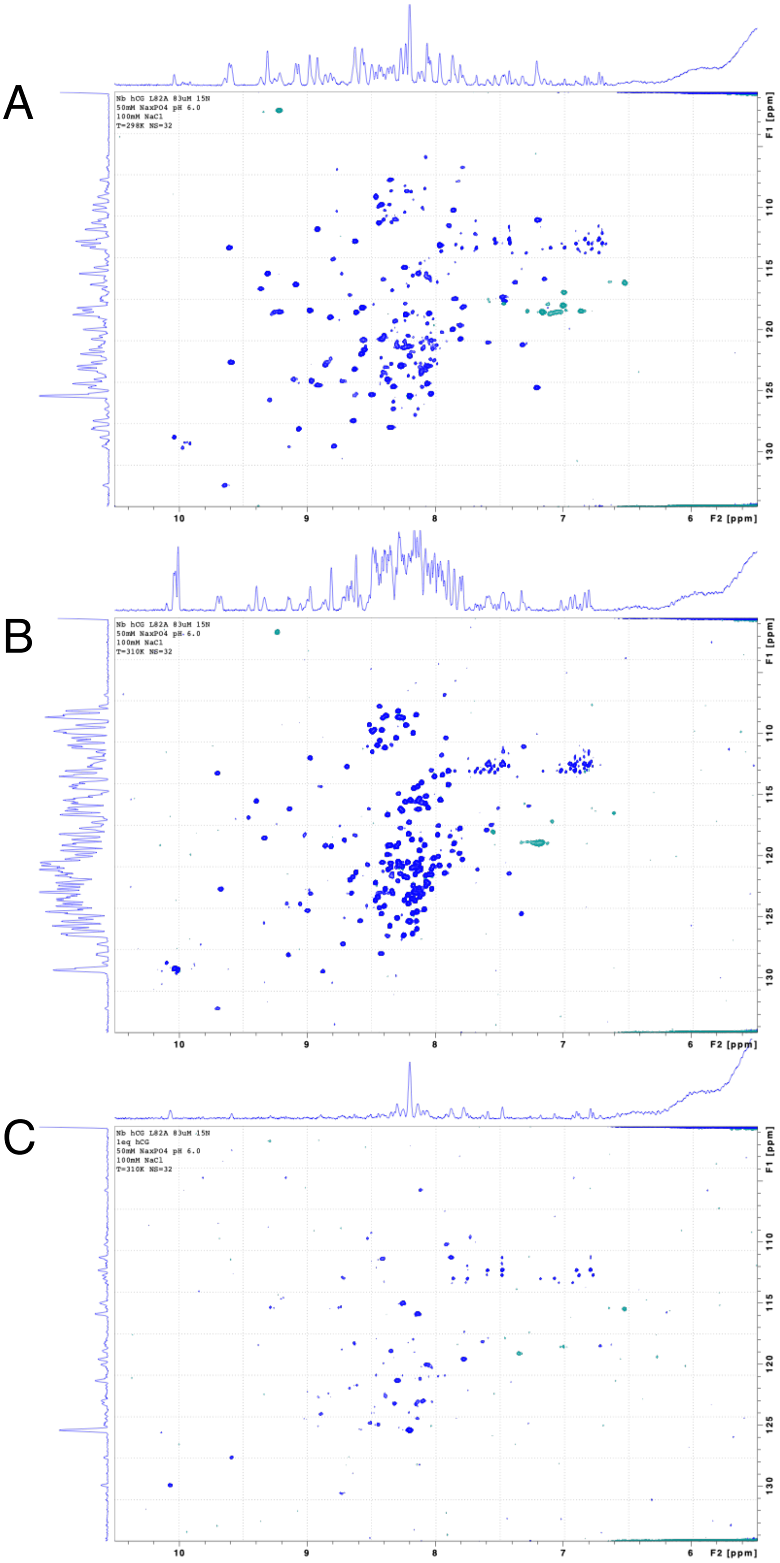


**Figure S2. Observing ligand-induced nanobody folding by NMR.** (**A**) $^{15}$N-HSQC of the anti-hCG nanobody L89A variant at 25 °C shows a well-dispersed spectrum, indicating a folded protein. (**B**) Increasing the temperature to 37 °C results in the appearance of a population of collapsed signals, corresponding to partial nanobody unfolding. (**C**) Addition of one equivalent of hCG results in the disappearance of the set of unfolded signals, signal broadening and loss of intensity, consistent with nanobody folding upon formation of a high molecular weight (~50 kDa) nanobody–hCG complex.

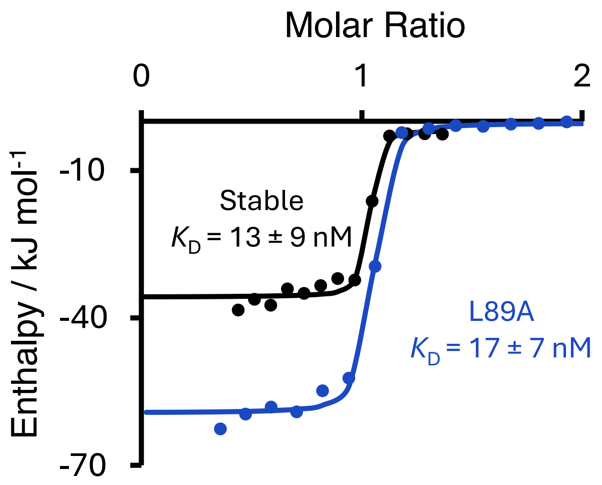


**Figure S3. Cavity-creating mutations in the nanobody framework does not affect binding affinity.** Isothermal titration calorimetry experiments at 25°C show similar affinities for both the stable nanobody and for the nanobody incorporating the destabilizing L89A mutation.

**Ligand-induced folding enthalpic fingerprint in calorimetric data:**

To calculate the unfolding enthalpy of the anti-hCG nanobody, we combined stability data from thermal and chemical denaturations to reconstruct the unfolding free energy-to-temperature dependence, following the Gibbs-Helmholtz equation (Eq. S1):

$$\Delta G(T) = \Delta H_{\mathrm{m}} \left(1 - {}^{T}/_{T_{\mathrm{m}}}\right) + \Delta C_{\mathrm{P}} \left[(T - T_{\mathrm{m}}) - T \ln\left({}^{T}/_{T_{\mathrm{m}}}\right)\right] \qquad \text{Eq. S1}$$

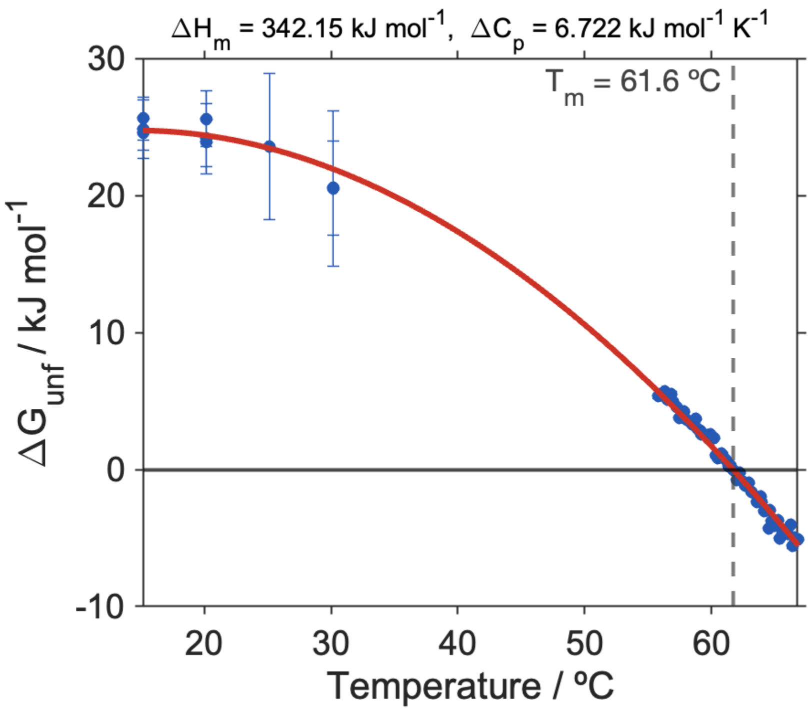


**Figure S4. Temperature-dependence of the stability of the anti-hCG nanobody.** We have measured nanobody stability at different temperatures, combining thermal and guanidinium-chloride denaturations and fitting the data to the Gibbs-Helmholtz equation to extract the change in enthalpy ($\Delta H_{\mathrm{m}}$) and heat capacity ($\Delta C_{\mathrm{P}}$) upon folding.

ITC measurements at 37 ºC show an enthalpy release 140 kJ mol$^{-1}$ higher for the anti-hCG L89A nanobody compared to the stable anti-hCG nanobody. We attribute this excess enthalpy to the contribution of ligand-induced folding. To evaluate this, we calculated the folding enthalpy at 37 ºC of the anti-hCG nanobody using the parameters obtained from the Gibbs-Helmholtz analysis (Eq. S2, Fig. S4):

$$\Delta H = \Delta H_{\mathrm{m}} + \Delta C_{\mathrm{P}}\,(T - T_{\mathrm{m}}) \qquad \text{Eq. S2}$$

This results in 177 kJ mol$^{-1}$. Given that the L89A variant is only 60% unfolded at 37 ºC, the expected enthalpic contribution from folding amounts to 107 kJ mol$^{-1}$, a value reasonably close to the 140 kJ mol$^{-1}$ we observe experimentally. Differences might arise due to entropy–enthalpy compensation mechanisms, and due to limitations in some of our assumptions. More specifically, we assume that the $\Delta H_{\mathrm{m}}$, $\Delta C_{\mathrm{P}}$, and $\Delta H_{\mathrm{Binding,\ 37\ ºC}}$ are the same for the stable nanobody and the L89A variant. Irrespective of these quantitative differences, this estimate validates that the large excess in enthalpy observed corresponds to the folding of the L89A variant.

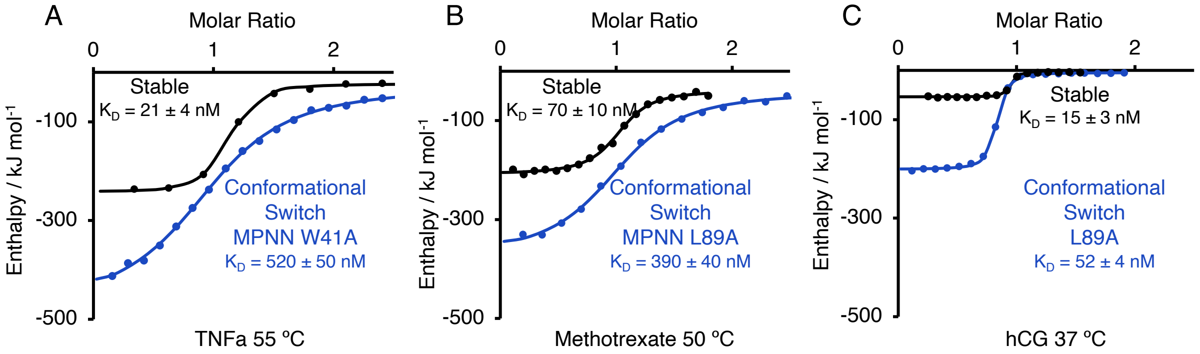


**Figure S5. The same framework-destabilization strategy generates conformational switches across different nanobodies scaffolds.** Isothermal titration calorimetry experiments indicate folding upon antigen binding of stable and destabilized variants of the anti-TNFa (**A**), anti-methotrexate (**B**) and anti-hCG (**C**) nanobodies. Increasing temperature shifts nanobody conformational equilibrium toward the unfolded state, activating the ligand-induced folding mechanism, as observed by the higher exothermic release upon binding. These mechanisms, however, introduce a trade-off that depends on the stability, reducing the apparent binding affinity compared to their stable variants at the same temperature.

**Experimental validation of the thermodynamic coupling of stability and affinity:**

Our thermodynamic model for ligand-induced folding establishes a dependence between the apparent affinity and stability (Fig. 1 and Eq. 1 in the main text). To evaluate whether our experimental results support this model, we measured binding affinities and stabilities for different destabilized variants of the three nanobodies studied, both at conditions at which they are stable and at which they undergo ligand-induced folding. For each variant and condition, we calculated the folding free energy ($\Delta G_f$) at the experimental temperature. Under destabilizing conditions—that is, at temperatures close to the melting temperature—thermal denaturation data directly provide folding free energies. For stable conditions, we calculated folding free energies by fitting the thermal denaturation data to the Gibbs-Helmholtz relationship (Eq. S1) using the $\Delta C_P$ and $\Delta H_m$ determined experimentally for the original nanobodies (see Fig. S4). We acknowledge that these could differ for the destabilized variants, which might introduce some uncertainty in the actual $\Delta G_F$ values, but our attempts at measuring chemical denaturations of the destabilized variants did not yield reliable stabilities. We note, however, that any differences would affect only the stabilities of the stable nanobodies and not the destabilized variants, and would thus not interfere with the validation of the ligand-induced folding model. Next, to determine the change in affinity, we performed ITC experiments for each variant both at 25 ºC and under destabilizing conditions (Fig. S5) to measure respectively the intrinsic ($K_{D,int}$) and apparent ($K_{D,app}$) dissociation constants. We then evaluate the relative change in affinity as the ratio $K_{D,app}$ / $K_{D,int}$, finding that it follows the predicted dependence on stability (Fig. 3 in the main text).

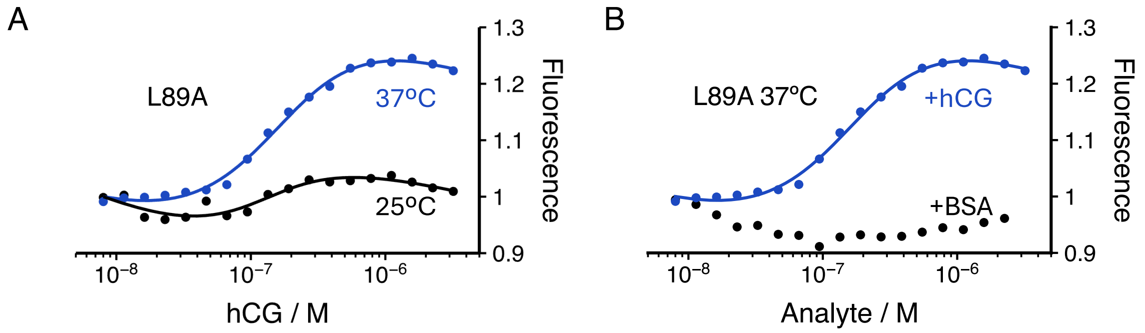


**Figure S6. Fluorescence signal transduction requires both nanobody destabilization and binding to the specific antigen.** Engineered anti-hCG nanobody L89A shows a significant signal change in buffer at 37 °C compared to 25 °C (**A**) and specific to hCG (**B**).

**Table S1. Thermostability data for the anti-hCG nanobody variants.**

| | Variant | Tm / °C | Reversibility / % |
|---|---|---|---|
| | Stable | 61 | 95 |
| Disulphide Removal | C22A/C92A | 40 | 90 |
| | C22A | 37 | 68 |
| | C22G | 34 | 80 |
| | C22G/C92M | 33 | 85 |
| | C22G/C92A | 33 | 77 |
| | C22S/C92S | 32 | 89 |
| | C22A/C92M | 32 | 75 |
| | C22G/C92W | 32 | 65 |
| | MPNN* C22S/C92S | 39 | 69 |
| Cavities | V87A | 58 | 91 |
| | I78A | 50 | 94 |
| | L21A | 41 | 90 |
| | L89A | 36 | 87 |
| | W41A | 27 | 82 |
| | MPNN* L89A | 41 | 85 |

* Variants based on a stabilized nanobody framework, which we previously reported on *Pejenaute et al. Prot Eng Des & Sel (2025) 38:gzaf010*.

**Table S2. Thermostability data for the anti-TNFa and anti-methotrexate nanobody variants.**

| | Variant | Tm / °C | Reversibility / % |
|---|---|---|---|
| TNFa | Stable | 66 | 56 |
| | L21A | 55 | 80 |
| | L89A | 49 | 58 |
| | W41A | 47 | 68 |
| | MPNN* L21A | 54 | 89 |
| | MPNN* L89A | 52 | 91 |
| | MPNN* W41A | 47 | 93 |
| | MPNN* C22S/C92S | 52 | 0 |
| Methotrexate | Stable | 69 | 72 |
| | MPNN* L21A | 56 | 72 |
| | MPNN* L89A | 50 | 80 |
| | MPNN* W41A | 47 | 91 |
| | MPNN* C22S/C92S | 46 | 48 |

* Variants based on an optimized nanobody framework, which we previously reported on *Pejenaute et al. Prot Eng Des & Sel (2025) 38:gzaf010*.

**Table S3. Amino acid sequences of the nanobodies employed in this study.**

| |
|---|
| **Human Chorionic Gonadotropin / 1HCV** |
| DVQLQESGGGLVQAGGSLRLSCAASGRTGSTYDMGWFRQAPGKERESVAAINWDSARTYYASSV<br>RGRFTISRDNAKKTVYLQMNSLKPEDTAVYTCGAGEGGTWDSWGQGTQVTVSS |
| **Tumor Necrosis Factor alpha / 5M2J** |
| QVQLVESGGGLVQPGGSLRLSCAASGFTFSNYWMYWVRQAPGKGLEWVSEINTNGLITKYPDSV<br>KGRFTISRDNAKNTLYLQMNSLSKPEDTALYYCARSPSGFNRGQGTQVTVSS |
| **Methotrexate / 3QXV** |
| QVQLVESGGGLVQAGGSLRLSCAASRRSSRSWAMAWFRQAPGKEREFVAKISGDGRLTTYGDSV<br>KGRFTISRDNAEYLVYLQMDSLKPEDTAVYYCAADDNYVTASWRSGPDYWGQGTQVTVSS |
